# Sixteen oogenesin genes are dispensable for fertility *What is the significance of the dispensability of genes expressed in germ-cell?*

**DOI:** 10.1101/2023.06.01.543189

**Authors:** Johan Castille, Dominique Thépot, Sophie Fouchécourt, Bruno Passet, Nathalie Daniel-Carlier, Jean-Luc Vilotte, Philippe Monget

## Abstract

Gene knockout experiments have shown that many genes are dispensable for a given biological function. The Oogenesin/Pramel family contains almost 85 paralogs, about thirty of which are specific to female (as well as male for some of them) germ cells. In this paper, we show that the deletion of a block of around 1Mb containing sixteen paralogous genes of the Oogenesin/Pramel family specific to germ cells, including Oogenesin-2, -3 and -4, has no consequences on fertility or prolificacy in mouse both sexes. The dispensability of these genes is probably due to the compensation by the other germ-cell specific paralogs.

## Introduction

The functional importance of gamete-expressed genes has become a crucial and acute issue with the availability of functional genomics tools. In the mouse, several genes of ubiquitous expression were targeted in the oocyte by using the cre-lox approach. It leads most of times to dramatic phenotypes such as a huge alteration of the reserve of primordial follicles (pTEN [1]) or the loss of oocytes with an early blockage of follicular growth (Omcg1 [2]; Mdm2 [3]). Several other genes, highly or exclusively expressed in the oocyte were invalidated by a general knock-out approach, leading to different phenotypes: sterility with a blockage of follicular growth at the primary follicles (GDF9: in the mouse [4], in the pig [5]), sterility with a huge defect of oocyte maturation (PATL2: [6]), sterility with an arrest of early embryonic development (Maternal genes Nlrp5, Stella, Nmp2, Zar1…[7]).

In males also, numerous knockout mouse models leading to male sterility were created in the past two decades (for reviews: [8][9][10]). Invalidation of germ cell specific genes allows revealing their role in spermatogenesis. According to the phenotype, the fertility impairment can occur at various stages of spermatogenesis. Some genes are involved in spermatogonia maintenance and/or differentiation, as RNA helicase *Ddx5* (DEAD box helicase 5), that has a multifaceted role within undifferentiated spermatogonia [11]. Other genes are involved in meiosis or chromatin organization, like HSF2BP, partner of the tumor suppressor BRCA2, whose inactivation results in male infertility due to a severe homolog recombination defect during meiosis [12]. Another group of genes are involved in sperm motility, like SLO3 [13]. Finally, a further group of genes are involved in fertilization, as Izumo and Juno [14].

In 2003, we have characterized one of the most expanding family of more than 85 genes in the mouse genome, the Oogenesin/Pramel family, whose more than thirty are predicted to be specifically expressed in male and/or female germ cells. One of these genes, called Oogenesin (Oog) that we renamed Oogenesin-1 (Oog1), had been first discovered the same year [15]. We have shown by RT-PCR and *in situ* hybridization that three others, that we called Oog2, -3 and -4, are exclusively expressed in murine oocytes from primary to large antral follicles, with a low expression level also observed in the testis for Oog4 [16]. This family has been extensively characterized since [17]. Oog1 is also expressed in two- and four-cell-stage embryos [15]. Moreover, it is a potential binding partner of the RAL guanine nucleotide dissociation stimulator (RALGDS), one of the RAS effector proteins [18]. The Oog/Pramel gene family is located on two main regions on chromosome 4 and 5 [16][17]. The *Oog*2, -3 and -4 are located on chromosome 4 between 143.1Mb and 143.9Mb (Table 2).

**Table 1.**
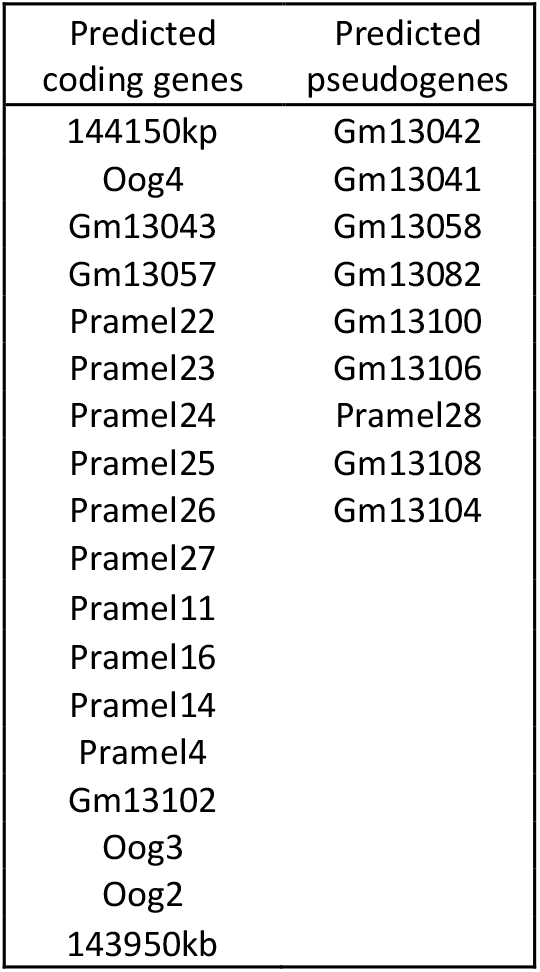
The sixteen Oogenesin/Pramel genes located on chromosome 4 between 144150kb and 143950kb. Nine pseudogenes from this family are also located in this region, between the predicted coding genes.

**Table 2.**
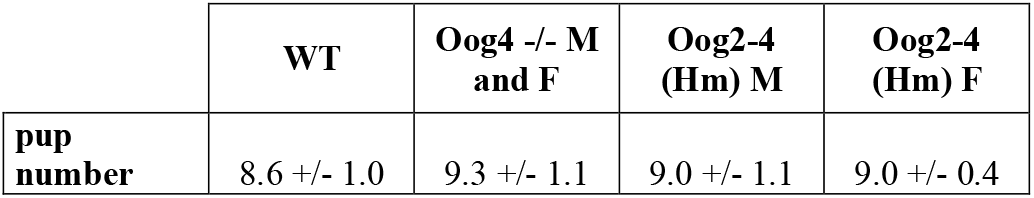
Prolificacy of Oog4 -/- and Oog2-4 mice (mean+/-SED; n = 6 to 12 crosses).

|  | WT | Oog4 <sup>-/-</sup> M and F | Oog2-4 (Hm) M | Oog2-4 (Hm) F |
| --- | --- | --- | --- | --- |
| pup number | 8.6 +/- 1.0 | 9.3 +/- 1.1 | 9.0 +/- 1.1 | 9.0 +/- 0.4 |

The gene encoding Oog2 has already been invalidated, with no effect on mouse fertility (https://www.mousephenotype.org/data/genes/MGI:2684035). In the present paper, we have invalidated the germ-cell specific gene *Oog4*, and the block of around 1Mb encompassed by *Oog2* and *Oog4*, containing thirteen other germ-cell specific *Oog* genes including *Oog3*.

## Materials & Methods

Animal ethical concerns were controlled and validated by the French *Ministère de l’enseignement supérieur, de la recherche et de l’innovation* and *Ministère de l’agriculture et de la souveraineté alimentaire*, under the authorization of the ethics committee COMETHEA. Generation of the *Oog4* knockout mice (COMETHEA authorization: APAFIS#28708-2020111711192750 v3) was achieved by deleting the gene transcription unit using two guides RNA (3’ *Oog4* and 5’ *Oog4*, Suppl. Table 1) that were electroporated into FVB/N mouse one-cell embryos alongside the Cas9 nuclease protein, as illustrated in Figure 1A and B and following already described protocols [19]. Resulting pups were screened by PCR analysis of their tail-derived genomic DNA using different set of primers (Suppl. Table 1 and 2). Transgenic mice were notably identified by the production of a PCR amplification fragment using primers Oog4AaF and Oog4PromF (see suppl. Tables 1 and 2). Sequence analysis of this amplified genomic DNA allowed confirming the precise deletion of the gene transcription unit (Suppl. Figure 1). Inter-crossing of transgenic mice was used to generate heterozygous or homozygous Oog4Del mice that were similarly identified by PCR screening (Suppl. Table 2).

**Figure 1.**
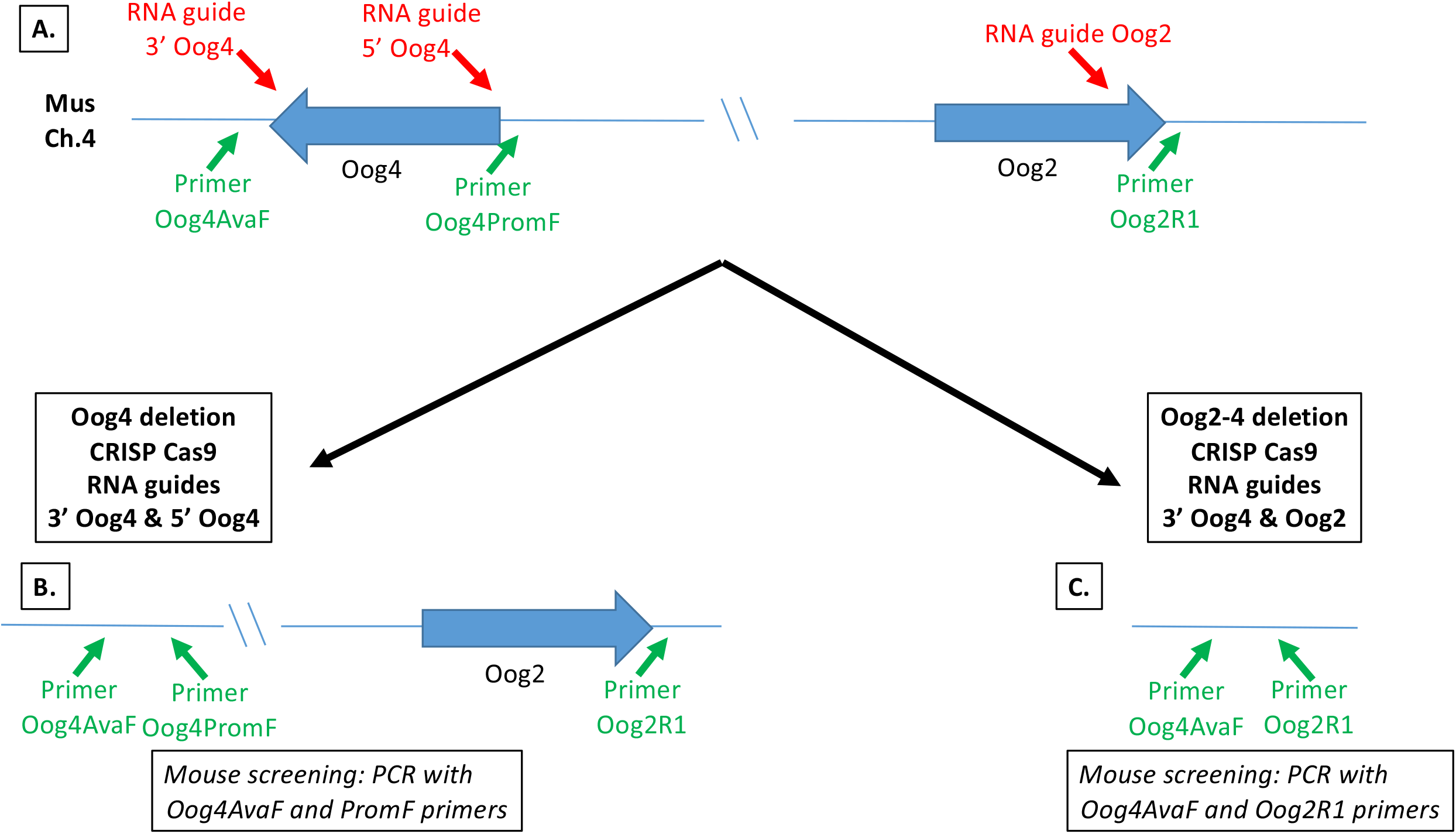
Deletions of Oogenesin 4 and of the Oogensin 4 – Oogenesin 2 gene cluster A. Schematic representation of the Oogenesin gene cluster on mouse chromosome 4 (Ch4). The Oogenesin gene cluster located on mouse chromosome 4 is schematically shown. The transcription units of the *Oog4* and *Oog2* genes are indicated as well as their orientations. The transcription unit of the *Oog4* gene extends from ∼ nucleotide 143176894 (5’) to nucleotide 143163734 (3’) and that of the *Oog2* gene from ∼ nucleotide 143917299 (5’) to nucleotide 143923504 (3’) according to GenBank NC_000070.7. Locations of the different RNA guides used to delete either *Oog4* or the entire gene cluster are indicated as well as those of the primers used to identify genome-edited mice. The figure is not at scale. B. Deletion of the *Oog4* transcription unit. The deletion was achieved through electroporation of mouse embryos at the pronuclei stage with the Cas9 protein (Alt-R S.p. Cas9 nuclease V3, Cat.1081058, Integrated DNA Technology) and a mixture of 3’ Oog4 and 5’ Oog4 RNA guides (Integrated DNA Technology), as indicated. The resulting mouse chromosome 4 Oogenesin gene cluster is schematically illustrated. Identification of genome-edited mice was performed by PCR amplification of the genomic DNA with the Oog4AvaF and Oog4PromF primers. Due to the length of the *Oog4* transcription unit and the 60s elongation time used in the PCR cycles, a DNA amplification could only be observed with DNA from genome-edited mice. C. Deletion of the *Oog4 – Oog2* gene cluster. The deletion was achieved through electroporation of mouse embryos at the pronuclei stage with the Cas9 protein (Alt-R S.p. Cas9 nuclease V3, Cat.1081058, Integrated DNA Technology) and a mixture of 3’ Oog4 and Oog2 RNA guides (Integrated DNA Technology), as indicated. The resulting mouse chromosome 4 lacking the Oogenesin gene cluster is schematically illustrated. Identification of genome-edited mice was performed by PCR amplification of the genomic DNA with the Oog4AvaF and Oog2R1 primers. Due to the length of the gene cluster and the 60s elongation time used in the PCR cycles, a DNA amplification could only be observed with DNA from genome-edited mice.

Similarly, generation of mice lacking the *Oog2-Oog4* gene cluster on chromosome 4 (COMETHEA authorization: APAFIS#28708-2020111711192750 v3) was achieved using two other guides RNA (3’ *Oog4* and *Oog2*, Suppl. Table 1) that were electroporated into FVB/N mouse one-cell embryos alongside the Cas9 nuclease protein, as illustrated in Figure 1A and C and following the same protocol. Resulting pups were screened by PCR analysis of their tail-derived genomic DNA using different set of primers (Suppl Tables 1 and 2). Transgenic mice were notably identified by the production of a PCR amplification fragment using primers Oog4AaF and Oog2R1 (see suppl. Tables 1 and 2). Sequence analysis of this amplified genomic DNA allowed confirming the precise deletion of the gene cluster (Suppl. Figure 2). Inter-crossing of transgenic mice was used to generate heterozygous or homozygous Oog2-4Del mice that were similarly identified by PCR screening (Suppl. Table 2).

### Results: Analysis of *Oog4* and *Oog2-Oog4* knockout mice

Two lines for which the *Oog4* gene has been invalidated were studied, the Oog4.1 and 4.6 lines. Males and females were fertile very quickly after breeding, crosses between homozygous (Hm) males and females with wild type counterpart giving an average litter size of 9.3 ± 1.1, which is not significantly different from control mice of the same FVB/N strain (8.6 ± 1.0; Table 1). In addition, the reproductive organs were weighed, without any difference with the control line (Figure 2).

**Figure 2.**
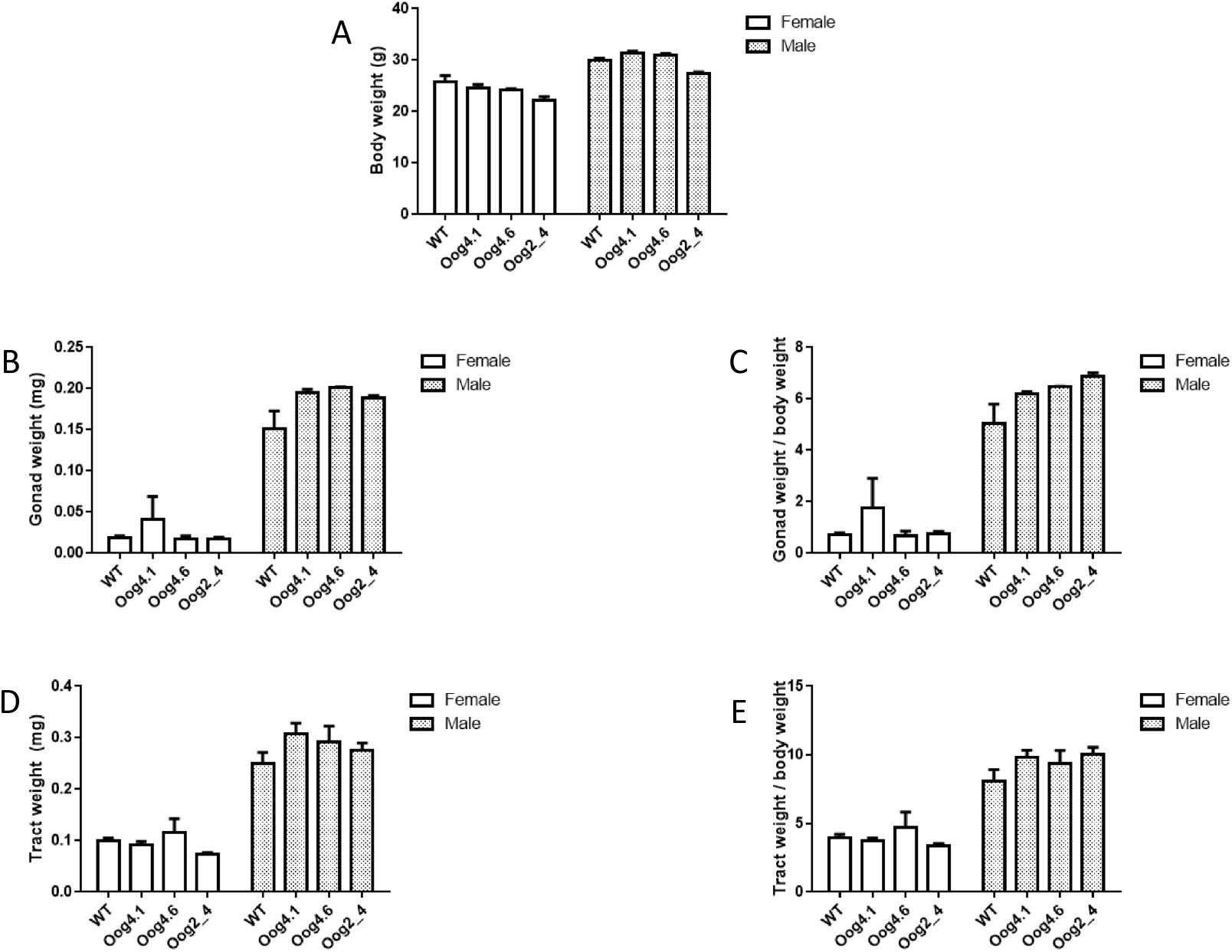
Weight of the body (A), of the gonads (B), and of the uterus and seminal vesicles (D) of WT mice, of both Oog4.1 and 4.6 Oog4-/-lines, and of the Oog2_4 female and male mice (animals in which the *Oog4 – Oog2* gene cluster has been deleted). C and E: ratio gonad weight/body weight (C) and ratio uterus or seminal vesicle/body weight (E) on the same animals.

We then decided to delete the block (*Oog2-Oog4*, approximately 1Mb) of Oog/Pramel genes located on chromosome 4 between *Oog2* and *Oog4*, containing *Oog2, Oog3, Oog4* and thirteen other paralogs predicted *in silico* to be specifically expressed in the ovary, and in the testis for some of them (Table 2). Very surprisingly, we observed no alteration in male and female fertility, nor in prolificacy (Table 1). No difference was observed in terms of the weight of the gonads and the female and male genital tracts were normal compared to controls (Figure 2). Our experience in phenotyping mice without any alteration of fertility or prolificacy led us to stop all experimentation on these animals, to avoid sacrificing many animals to show that the histology of the ovaries and testicles was normal, and that the levels of the main reproductive hormones were similar to control mice.

## Discussion

### Existence of paralogs that compensate the loss of genes?

First of all, concerning the Oog/Pramel gene family, there are still at least thirty germ cell-specific paralogs, in particular in a locus on chromosome 5, which may be able to compensate the loss of the thirteen genes deleted here, the other genes of the family being of ubiquitous expression. To test this hypothesis, it would be necessary to invalidate all these genes one by one, which is very costly in terms of finances and animal lives.

The presence of other paralogs is probably not sufficient to explain the dispensability of some paralogs. Indeed, among the Nlrp5 (Mater) family of oocyte genes, i.e. *Nlrp4e, Nlrp9a, Nlrp9b, Nlrp9c*, the *in vivo* invalidation of *Nlrp5* renders female mice sterile due to embryonic mortality at stage 2 cells [20], whereas the invalidation of the three *Nalp9a/b/c* together has no phenotypic consequences [21]. Furthermore, the *in vitro* invalidation of *Nalp4e* results in the arrest of embryo development between the 2- and 4-cell stage [22]. So, in this oocyte-specific Nlrp gene family, *Nlrp5* is crucial for fertility (and seems to be for *Nlrp4e*), whereas the others seem useless. Another example comes from the three members of the Oosp family of oocyte-specific genes (*Oosp1*, -2 and *-3* [23]): none appears to be essential for fertility [24], even if the prolificacy of Oosp triple knockout (KO) females is reduced, suggesting a reduction of embryo viability.

### Evolution of paralogs by gene conversion or positive selection to predict dispensability?

In eukaryotes, genes can evolve, among other things, by birth, by pseudogenization, by positive selection, by negative (purifying) selection, by gene conversion, and/or by duplication [25]. In particular if the duplications are recent and fairly massive, they can be followed by negative selection or gene conversion, molecular phenomena which are likely to, respectively, conserve or homogenize the duplicated sequences. In these cases, one can hypothesize that the newly formed paralogous genes show a certain functional redundancy, with the consequence of an absence of phenotype when one of the paralogs is mutated (one cannot know if it is linked to a total loss of function), because another non-mutated paralog takes over [26]. In other cases, recent or more ancient duplications are followed by the evolution of paralogs by positive selection which, in contrast to negative selection or gene conversion, can lead to functional molecular diversification of the different paralogs that can acquire new specific biological roles (neofunctionalization). In these cases, the general hypothesis is that the loss of function of one of them could not be compensated by its paralogs. For example, among the paralogs specifically expressed in the murine oocyte that we have identified [27], individual invalidation of Nlrp5 and Nlrp4e, leads in both cases to a drastic phenotype of sterility with early embryonic death [20][22]. In my lab, we have shown that this gene family has been subjected to positive selection [28]. To our knowledge, this is one of the rare known examples of potential link between positive selection and absence of functional compensation in vertebrates.

Interestingly, some members of the Oog/Pramel family are subjected to positive selection [17], suggesting a functional diversification of these genes, which is contradictory with the absence of phenotypic consequences on fertility of the 1Mb deletion within the *Oog2-Oog4* genomic region. Further investigations may hopefully help to understand this contradiction.

### Necessity of a deep phenotyping study of mutant animals?

“Absence of evidence does not mean evidence of absence” said Carl Sagan. In our case, that means that it is possible that we could (or not) find a subtle phenotype in animals lacking the sixteen *Oog* genes such as less primordial follicles (because of an impairment in the constitution of the pool of the ovarian reserve), accelerated ovarian aging with early menopause as in *AMH*^-/-^ mice [29], impaired recruitment of ovarian follicles due to disturbed functioning of tyrosine kinase signaling pathway (linked to RAS), etc…

More generally, it is also possible that some of the known dispensable genes prove to be crucial only under extreme experimental conditions. For example, embryonic stem cells from *Zsacn5b* KO females showed an impaired DNA damage response after irradiation, compared to controls [30]. Moreover, genetic background may influence the dispensability of genes. This has been demonstrated for *Usp26* KO males, which are fertile on a C57Bl6/DBA2 background, and subfertile after several backcross on the DBA2 background [31]. Testing this hypothesis would require systematic crosses over several generations and in several genetic backgrounds. Of note, our *Oog4* and *Oog2-4* deletions were generated on a FVB/N genetic background, whereas the *Oog2* deletion was generated on a C57/Bl6 genetic background (https://www.mousephenotype.org/data/genes/MGI:2684035).

Whatever the case, we thoroughly phenotyped in our lab two lines of invalidated mice (data not published: IGF1R KO) because we speculated a subtle impaired ovarian function, although their fertility and prolificacy were normal. For this purpose, we sacrificed almost 1000 mice, and we found nothing. All this raises important ethical issues for us and allowed us to be reasonable in the phenotyping of Oog2-Oog4 deleted mice.

### Duplications “for nothing” of germ cells specific genes?

In conclusion, we ask the question of the significance of the existence of duplicated Oogenesin for reproduction. Could it be that some of the massive germ cell-specific gene duplications have no physiological nor evolutive significance? And that these types of tissue-specific duplications also exist de novo in other somatic tissues, but that these duplications are not seen in the databases because they are not transmitted to offspring?

In conclusion, we ask the question of the significance of the existence of these dispensable genes for a biological function. From an evolutionary point of view, there are functions which gradually become unnecessary, the outcome of this relaxation of function being the loss of genes by pseudogenization, such as the vitellogenin genes in viviparous species, or that of the GULO gene (necessary for the synthesis of vitamin C) in primates and bats. In this case, these germ-cell specific genes should be in the process of pseudogenization, as is the case of the nine pseudogenes of the oogenesin family of the *Oog2-4* locus that we have deleted (Table 2). We described the existence of such pseudogenes of the germ cell specific Nlrp family [28], which argues in favor of a general expansion of these germ-cell families followed by a relaxation of function, preliminary to a progressive loss of genes. Another question would be whether there are more dispensable genes among germ cell specific genes than among genes of somatic expression. If so, maybe that would have given Ken Mc Natty the opportunity to say that “germ cell-specific genes are a little crazy” [32].

## Acknowledgments

We thank Clément Gérard for its help

