## Supplemental Figures and Tables for "Sixteen oogenesin genes are dispensable for fertility *What is the significance of the dispensability of genes expressed in germ-cell?*"

**Suppl. Table 1. Sequences of the primers and RNA guides.**

| RNA guide or Primer name | Sequence (5'-3') | Alignment on mouse chromosome 4 (GenBank NC_000070.7) |
|---|---|---|
| RNA guide 3' Oog4 | AGTATCAGGAGCATAGACCTGGG | 143158335-143158357 |
| RNA guide 5' Oog4 | TCCACCCATTCAATCTTATTTGG | 143180419- 143180441 |
| RNA guide Oog2 | TTGGTTACTTCAACAATACTAGG | 143919671- 143919649 |
| Primer Oog4AvaF | TCTGCTTCTGGGAAATACTGTGTC | 143158141- 143158164 |
| Primer Oog4AvaR | TCTTGAAGGAGATGACTTGCTCTG | 143158636- 143158613 |
| Primer Oog4PromF | GTAGGGAATCACAGAAGGAAGG | 143180725- 143180704 |
| Primer Oog4PromR | CACTTGAAGAGACAGACTCCAG | 143180186- 143180207 |
| Primer Oog2F | TGTAGTCTCTAAGTTGGGTGG | 143919334- 143919354 |
| Primer Oog2R1 | GAGGTCTAAATAGAGTGGTTAAAG | 143919835- 143919812 |
| Primer Oog4IntF | CTGGAGCGATCATTTTGTGCAA | 143166462- 143166483 |
| Primer Oog4IntR | GAATGTATCTTTCGAAGAAGACA | 143166773- 143166751 |

**Suppl. Table 2. Mouse PCR genotyping.** PCR conditions were 94°C/3min, 40x(94°C/30s ; 60°C/30s ; 72°C/30s), 72°C/min. Primer sequences are listed in Suppl. Table 1.
-: no PCR amplification signal. +: PCR amplification signal.

| Mouse genotype/PCR primer set | Oog4AvaF/ Oog4R | Oog4PromF/ Oog4PromR | Oog2F/ Oog2R1 | Oog4IntF/ Oog4IntR | Oog4PromF/ Oog4AvaF | Oog4AvaF/ Oog2R1 |
|---|---|---|---|---|---|---|
| WT | + | + | + | + | - | - |
| *Oog4*$^{+/Del}$ | + | + | + | + | + | - |
| *Oog4*$^{Del/Del}$ | - | - | + | - | + | - |
| *Oog2-4*$^{+/Del}$ | + | + | + | + | - | + |
| *Oog2-4*$^{Del/Del}$ | - | - | - | - | - | + |

```
Query  20         TTTATGAAGTGATCTCTAAGGACCCCAGAACAAGTCATCTCCTCGAAGACAAAGCCCTAG  79
                  |||||||||||||||||||||||||||||||||||||||||||| |||||||||||||||
Sbjct  143158188  TTTATGAAGTGATCTCTAAGGACCCCAGAACAAGTCATCTCCTCAAAGACAAAGCCCTAG  143158247

Query  80         ATCAGTCTCCTGCTGGGAGCAGGAACACAGTGCTGGGGCAGCAGTGGGTCCAGAGCTGGG  139
                  ||||||||||||||||||||||||||||||||||||||||||||||||| ||||||||||
Sbjct  143158248  ATCAGTCTCCTGCTGGGAGCAGGAACACAGTGCTGGGGCAGCAGTGGGTACAGAGCTGGG  143158307

Query  140        ACTAGGGTATGAGGAGCATAGCCCTGGAGTATCAGGAGCATAGA  183
                  ||| ||||||||||||||||||||||||||||||||||||||||
Sbjct  143158308  ACTGGGGTATGAGGAGCATAGCCCTGGAGTATCAGGAGCATAGA  143158351


Query  184        ATTTGGGACACCTTTATCAATATTCCTTGAACCAAATATCTCTGCACGAAAGAAGGATAA  243
                  ||||||||||||||||||||||||||||||||||||||||||||||| ||||||||||||
Sbjct  143180436  ATTTGGGACACCTTTATCAATATTCCTTGAACCAAATATCTCTGCACAAAAGAAGGATAA  143180495

Query  244        AAAACTGAATATGACCAGAAAAATTCCTGACCGTGGGATGAATATCAGATGTCCATGAAG  303
                  |||||||||||||||||||||||||||||||| |||||||||||||||||||||||||||
Sbjct  143180496  AAAACTGAATATGACCAGAAAAATTCCTGACCATGGGATGAATATCAGATGTCCATGAAG  143180555

Query  304        CACTGACTTTACAACTCTGAGTTCATTTAtttttttAATGTATTCAGTTATTTGTTTATC  363
                  ||||||||||||||||||||||||||||||||||||||||||||||||||||||||||||
Sbjct  143180556  CACTGACTTTACAACTCTGAGTTCATTTATTTTTTTAATGTATTCAGTTATTTGTTTATC  143180615

Query  364        TTTTTCAGCTGGCTTGATATTTGCTTCTGTAACAGTCTTACTCTGTAGATAATCTTAACT  423
                  ||||||||||||||||||||||||||||||||||||||||||||||||||||||||||||
Sbjct  143180616  TTTTTCAGCTGGCTTGATATTTGCTTCTGTAACAGTCTTACTCTGTAGATAATCTTAACT  143180675

Query  424        AGGTTTCTCTGGAGTCCTTTTGCTACATCCTTCCTTCTG-GATTTCC  469
                  ||||||||||||||||||||||||||||||||||||||| |||| ||
Sbjct  143180676  AGGTTTCTCTGGAGTCCTTTTGCTACATCCTTCCTTCTGTGATTCCC  143180722
```

**Suppl. Figure 1. Sequence alignment of the amplified PCR genomic fragment using primers Oog4AvaF and Oog4PromF on the mouse genome: allele *Oog4*$^{Del}$.**

The amplified genomic DNA fragment obtained using primers Oog4AvaF and Oog4PrmF was aligned on the mouse genome by Blast (https://blast.ncbi.nlm.nih.gov/Blast.cgi). The best alignment was obtained on mouse chromosome 4 (GenBank NC_000070.7) as illustrated here, with two non-contiguous regions of homology, illustrating genomic DNA cutting by Cas9 nuclease in the regions targeted by the guide RNAs (3’ Oog4 and 5’ Oog4) and DNA religation resulting in deletion of the *Oog4* gene transcription unit.

```
Query  9          GGG-CCTACCATTTTATGAAGTGATCTCTAAGGACCCCAGAACAAGTCATCTCCTCGAAG  67
                  ||| |||||||||||||||||||||||||||||||||||||||||||||||||||| |||
Sbjct  143158176  GGGCCCTACCATTTTATGAAGTGATCTCTAAGGACCCCAGAACAAGTCATCTCCTCAAAG  143158235

Query  68         ACAAAGCCCTAGATCAGTCTCCTGCTGGGAGCAGGAACACAGTGCTGGGGCAGCAGTGGG  127
                  ||||||||||||||||||||||||||||||||||||||||||||||||||||||||||||
Sbjct  143158236  ACAAAGCCCTAGATCAGTCTCCTGCTGGGAGCAGGAACACAGTGCTGGGGCAGCAGTGGG  143158295

Query  128        TCCAGAGCTGGGACTAGGGTATGAGGAGCATAGCCCTGGAGTATCAGGAGCATAGA  183
                  | ||||||||||||| ||||||||||||||||||||||||||||||||||||||||
Sbjct  143158296  TACAGAGCTGGGACTGGGGTATGAGGAGCATAGCCCTGGAGTATCAGGAGCATAGA  143158351


Query  180        TAGAATTGTTGAAGTAACCAATCATGTTATTGAACTAATGAGTCTCTTGACTTTAGCATG  239
                  ||| ||||||||||||||||||||||||||||||||||||||||||||||||||||||||
Sbjct  143919651  TAGTATTGTTGAAGTAACCAATCATGTTATTGAACTAATGAGTCTCTTGACTTTAGCATG  143919710

Query  240        TGAATTATAATCTCAAACTCATAGTCTCTTCTTTAAGGATGCTATATAAAGAGTTAGAGA  299
                  ||||||||||||||||||||||||||||||||||||||||||||||||||||||||||||
Sbjct  143919711  TGAATTATAATCTCAAACTCATAGTCTCTTCTTTAAGGATGCTATATAAAGAGTTAGAGA  143919770

Query  300        CCATGTTAATGGCTACTGAAGAATTACTGACTTTAGTGTTTCTTTAACCACTCTATTTAG  359
                  ||||||||||||||||||||| ||||||||||||||||||||||||||||||||||||||
Sbjct  143919771  CCATGTTAATGGCTACTGAAGGATTACTGACTTTAGTGTTTCTTTAACCACTCTATTTAG  143919830

Query  360        ACCTCAA  366
                  |||||||
Sbjct  143919831  ACCTCAA  143919837
```

**Suppl. Figure 2. Sequence alignment of the amplified PCR genomic fragment using primers Oog4AvaF and Oog2R1 on the mouse genome: allele *Oog2-4*$^{Del}$.**

The amplified genomic DNA fragment obtained using primers Oog4AvaF and Oog2R1 was aligned on the mouse genome by Blast (https://blast.ncbi.nlm.nih.gov/Blast.cgi). The best alignment was obtained on mouse chromosome 4 (GenBank NC_000070.7) as illustrated here, with two non-contiguous regions of homology, illustrating genomic DNA cutting by Cas9 nuclease in the regions targeted by the guide RNAs (3' Oog4 and Oog2) and DNA religation resulting in deletion of the Oogenesin gene cluster.
